# Vector-valued dopamine improves learning of continuous outputs in the striatum

**DOI:** 10.1101/2022.11.30.518587

**Authors:** Emil Wärnberg, Arvind Kumar

## Abstract

It is well established that midbrain dopaminergic neurons support reinforcement learning (RL) in the basal ganglia by transmitting a reward prediction error (RPE) to the striatum. In particular, different computational models and experiments have shown that a striatum-wide RPE signal can support RL over a small discrete set of actions (e.g. no/no-go, choose left/right). However, there is accumulating evidence that the basal ganglia functions not as a selector between predefined actions, but rather as a dynamical system with graded, continuous outputs. To reconcile this view with RL, there is a need to explain how dopamine could support learning of dynamic outputs, rather than discrete action values.

Inspired by the recent observations that besides RPE, the firing rates of midbrain dopaminergic neurons correlate with motor and cognitive variables, we propose a model in which dopamine signal in the striatum carries a vector-valued error feedback signal (a loss gradient) instead of a homogeneous scalar error (a loss). Using a recurrent network model of the basal ganglia, we show that such a vector-valued feedback signal results in an increased capacity to learn a multidimensional series of real-valued outputs. The corticostriatal plasticity rule we employed is based on Random Feedback Learning Online learning and is a fully local, “three-factor” product of the presynaptic firing rate, a post-synaptic factor and the unique dopamine concentration perceived by each striatal neuron. Crucially, we demonstrate that under this plasticity rule, the improvement in learning does not require precise nigrostriatal synapses, but is compatible with random placement of varicosities and diffuse volume transmission of dopamine.

## Introduction

The basal ganglia are thought to be the main locus of reinforcement learning (RL) in the brain (Niv and Langdon, 2016). In particular, dopamine modulated long-term plasticity in the corticostriatal synapses is crucial for learning and fine-tuning skilled movements based on environmental feedback (Perrin and Venance, 2019). Combined with the striking observation that midbrain dopaminergic cells transmit a reward prediction error (RPE) to the striatum (Schultz et al., 1997; Kim et al., 2020), this has inspired a plethora of computational models of the basal ganglia implementing various forms of RL. Notably however, virtually all these models assume the set of actions that can be selected — the *action space* in RL terminology — is small and discrete (see e.g.Humphries et al., 2006; Stewart et al., 2012; Berthet et al., 2012; Bolado-Gomez and Gurney, 2013; Gurney et al., 2015; Baston and Ursino, 2015; Bogacz et al., 2016; Berthet et al., 2016; Dunovan et al., 2019; Bahuguna et al., 2019; Pozzi et al., 2020). Practically, this means that each action can be exclusively represented by a disjoint group of striatal neurons, sometimes called *action channels* (Redgrave et al., 1999). At their core, in each of these models there is some sort of competition between the action channels so that the selected action (or likely selected in probabilistic models) corresponds to the channel with the highest activity. This is consistent with a global RPE transmitted by dopamine that reinforces or depresses the corticostriatal synapses of the active channel.

However, there is now accumulating evidence that action space of the basal ganglia is not small and discrete, but rather multidimensional and continuous (Yin, 2014; Barter et al., 2015; Rueda-Orozco and Robbe, 2015; Park et al., 2020; Dhawale et al., 2021). For a multidimensional output, a global RPE is not as effective at driving learning as it is for discrete action channels. For example, a mouse learning to reach for a food pellet may need to learn to control its paw in the *x, y*, and *z* directions. Intuitively, a single error, perhaps proportional to the final distance to the target, would be less efficient than having a three-dimensional error signal representing the error in the three directions. More formally, producing a continuous and multidimensional output requires the basal ganglia to learn a function approximation rather than tabular values (Sutton et al., 1999). In this formulation, a scalar RPE signal transmitted to the entire network corresponds to perturbation learning (Chen and Goldberg, 2020), which is well-known to be considerably less efficient than vector valued feedback, especially for a network with multiple layers (Lillicrap et al., 2020).

Therefore, we asked if it would be possible for dopamine to support function approximation learning in the basal ganglia by carrying a vector-valued feedback signal from the midbrain back to the striatum. Such a feedback signal would manifest in the VTA and SNc in terms of cell tunings to various task-related variables, consistent with recent observations (Fan et al., 2012; Howe and Dombeck, 2016; Engelhard et al., 2019). However, one apparent problem with dopamine transmitting a vector-valued error is that dopaminergic axons do not precisely target specific neurons in the striatum, but instead release dopamine from a large number of varicosities that can then diffuse over a short distance through extracellular space (Arbuthnott and Wickens, 2007), thereby mixing any individual error components. In principle one could imagine this problem being solved by representing each action dimension in a spatially compact region that receives its private dopamine channel (Hamid et al. (2021) and Lee et al. (2022) have proposed similar ideas). Firstly however, experimental evidence suggests both individual striatal projection neurons (Rueda-Orozco and Robbe, 2015; Klaus et al., 2017; Weglage et al., 2021) and individual dopaminergic neurons (Avvisati et al., 2022) have mixed tuning rather than responding to a single task variable, as would be predicted from a naive parallel organization. Secondly, although there is a coarse-grain somatotopic organisation of the striatum (Hunnicutt et al., 2016) as well as the substantia nigra (Foster et al., 2021), the axonal arborizations of individual SNc neurons are huge and cover large portions of the striatum (up to 5%; Matsuda et al., 2009). Therefore, a large set of isolated parallel channels without cross-talk appears unlikely.

In this work, rather than separating the entire basal ganglia into fine-grained parallel channels, we propose that the mixing of multiple error components can be undone downstream from the striatum. In particular, we propose that if the striatofugal projections were to be subject to similar systematic long-term plasticity as the corticostriatal projections, then we can make use of *feedback alignment* (Lillicrap et al., 2016) to have striatum learn continuous outputs efficiently. We demonstrate that learning in a recurrent neural network model of the basal ganglia is better with a stylized model of diffuse mixing of dopaminergic feedback compared to a model where there is a homogeneous, scalar dopamine concentration in the entire striatum. Thus, we connect two seemingly unrelated observations: heterogeneous dopamine response and the involvement of striatum in learning continuous actions.

## Results

### Network model

As a model of the basal ganglia learning a skilled movement (such as an animal reaching for a food pellet or pressing a lever), we constructed a task wherein a recurrent neural network must learn to repeatedly output a given trajectory in *d*-dimensional space. The output trajectory is defined as the activity of *d* readout neurons. In our idealized model of a small piece of basal ganglia, we take the readout population to be the either the internal globus pallidus (GPi) or the substantia nigra pars reticulata (SNr) (see Barter et al., 2015, for experimental support).

This striatum projects to the readout population (GPi/SNr) and receives excitatory inputs from two input populations: cortex and thalamus (Fig. 1A). The task of the network is to adjust the input and recurrent synaptic weights in the striatum so that the readout matches the desired *d*-dimensional target *T*(*t*) as closely as possible (Fig. 1I).

**Figure 1:**
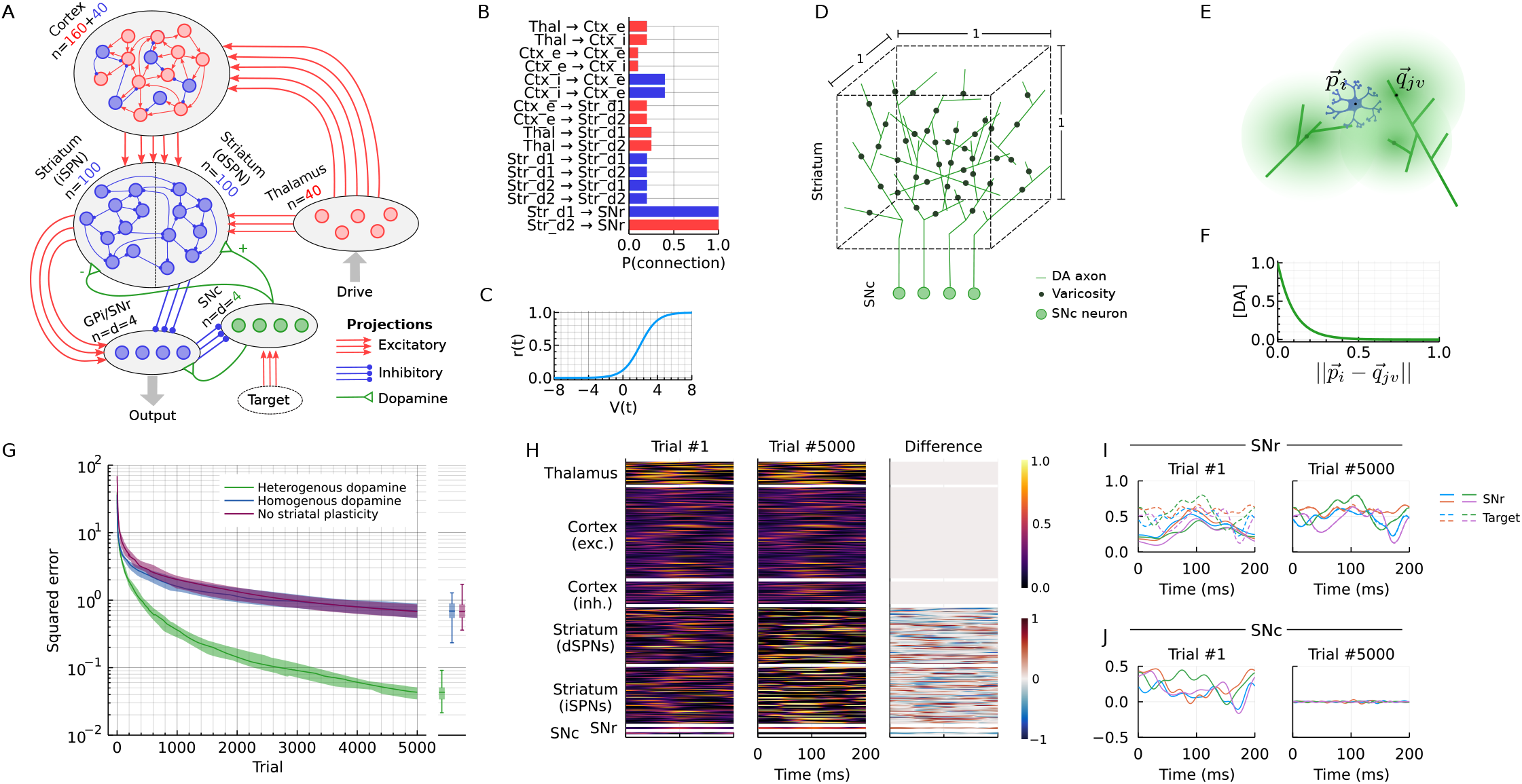
Heterogeneous dopamine improves learning in a RNN. (A) Network architecture. *n* indicates the number of units in each population. (B) Connection probability of a synapse for any pair of neurons in the network model. (C) Transfer function of each unit. D) Illustration of placement of varicosities in a unit cube. (E) Illustration of a striatal projection neurons (SPN) (located at 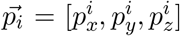) receiving a mix of dopamine from three close-by varicosities. (F) Relative dopamine concentration as a function of distance from the varicosity. (G) Convergence of the loss (L2) over trials for the model (heterogeneous dopamine) as well as two control models (homogeneous dopamine and no striatal plasticity). Each trial is one presentation of the pattern. Solid lines indicate the median of 25 runs; shaded areas show first and third quartiles. Boxplots show median, quartiles and min/max of the same 25 runs at the final (5000th) training trial. (H) Example activity in the network in the first and last trials of training. (I) Comparison of the 4-dimensional readout (i.e. SNr/GPi activity) with the target in the first and last trial of training. (J) The difference between the readout and the target (i.e. SNc activity) in the first and last trial of training.

In cortex and striatum, we model the sub-threshold membrane potential *V_i_*(*t*) of the *i*th neuron (or small group of neurons) as

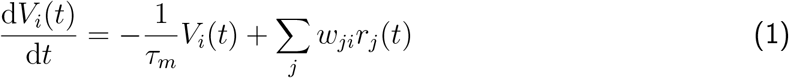

where *τ_m_* is the membrane time constant, *w_ji_* is the signed synaptic weight from neuron *i* to neuron *j* and *r_j_*(*t*) is the firing rate of unit *j* given as:

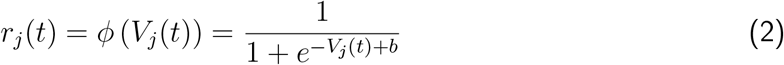

where the term *b* = 2 shifts the sigmoid to the right so that the firing rates are sparser when the inputs are balanced and the membrane potentials fluctuate around 0 (Fig. 1C).

The purpose of our model is to demonstrate that mixing of dopamine is not detrimental to vector-valued feedback, not to capture every detail of basal ganglia. Nevertheless, to make sure the learning set-up is fair, we added a number of constraints to the model. First, all connections except the readout are sparse, i.e. only a fraction of pairs of neurons are allowed to connect (Fig. 1B). Second, we required the signs of the weights *w_ij_* to match the sign of the projection (excitatory or inhibitory) throughout learning (Fig. 1A-B). For simplicity, we omitted the external globus pallidus (GPe) and subthalamic nucleus (STN) and modelled the indirect pathway as a direct excitatory projection from striatum to GPi/SNr. The sign of dopamine-driven plasticity was reversed for the striatal projection neurons in the indirect pathway (iSPNs; Fig. 1A). Because of our focus on dopamine-dependent learning in the striatum, we only include the cortex as a reservoir of rich but task-aligned dynamics, and do not consider any learning that might take place in the cortex itself. However, we include dopamine-dependent plasticity in all basal ganglia synapses: corticostriatal, thalamostriatal, striatostriatal and striatofugal.

### Derivation of the synaptic plasticity rule

To construct normatively appropriate learning rules for the plastic synapses, we note the task is to minimize the loss

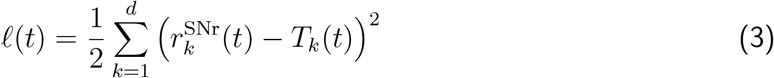

For the error to decrease over time, we would like the plasticity rule to change the weight of each corticostriatal synapse 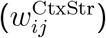 such that

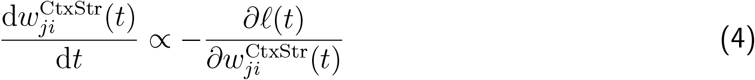

In Supplementary Text, we show that by expanding this partial derivative with two simplifications, only considering *ℓ* only at *t* (i.e. not backwards or forward in time) and treating the firing rates of other SPNs as fixed (Murray, 2019), we arrive at the following plasticity rule:

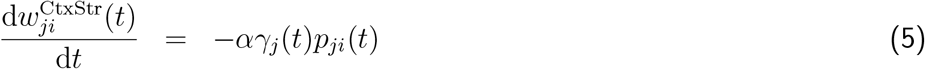

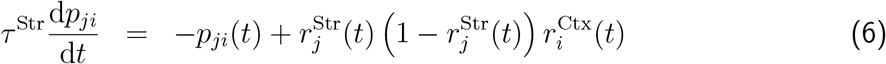

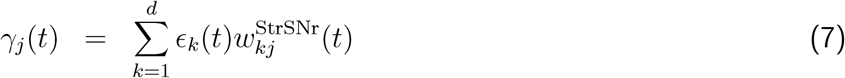

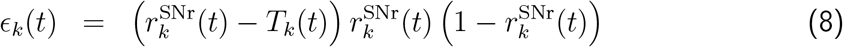

The plasticity rules for the thalamostriatal and striatostrial synaptic weights are fully analogous.

We interpret this plasticity rule in biological terms as follows. From Eq. 5, we see that the weight update depends on a neuron-specific factor *γ_j_* and a synapse-specific factor *p_ji_*. The latter is a low-pass filtered trace of a Hebbian-like product between pre- and postsynaptic firing. This could be identified as an eligibility trace (Izhikevich, 2007; Gurney et al., 2015) and we note it could be represented for example by the local concentration of calcium in the spine.

The eligibility trace *p_ji_* is multiplied by a “third factor” *γ_j_*. Experimental results suggest plasticity in corticostriatal synapses depends on three factors: presynaptic activity, postsynaptic activity, and dopamine (Fisher et al., 2017; Gerstner et al., 2018). Given that the two former are captured by *p_ji_*, we would like to associate *γ_j_* in Eq. 5 with dopamine. If we assume the number of dopaminergic neurons to be the same as the number of read-out neurons, we can assign *ε_k_* to the firing rate 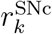. Note that we can handle negative values of *ε_k_* by loosely interpreting 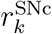 as the deviation from some baseline firing rate. This leaves just one problem: the coefficients used to sum to contribution of the dopaminergic cells in Eq. 7 should be the *downstream* striatofugal weights 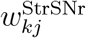, which are not available to the corticostriatal synapses. However, Murray (2019) showed that, if the readout weights 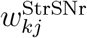 themselves are plastic, we can replace 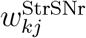 in Eq. 7 with a random value at only a minor cost to the convergence of the loss, thanks to a phenomenon called *feedback alignment* (Lillicrap et al., 2016). Therefore, we next asked whether the dopaminergic nigrostriatal projection could form such a random feedback matrix.

### Dopamine diffusion

To investigate if dopaminergic feedback could be used to communicate the third factor needed for the corticostriatal plasticity, we set up a stylized model of dopamine diffusion. We assumed each striatal neuron had a position 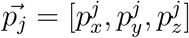 where 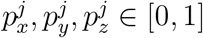. Second, we assumed that each SNc neuron sent axonal projections that covered the entire cube, and that axonal arbor of each SNc neuron has *N*^var^ = 10 varicosities randomly placed in the unit cube (Fig. 1D). Third, we assumed the dopamine released from each varicosity is proportional to the firing rate at the soma in the SNc and that the dopamine concentration decreases exponentially with distance from the varicosity. This gives the dopamine concentration *C_j_*(*t*) at striatal neuron *j* as

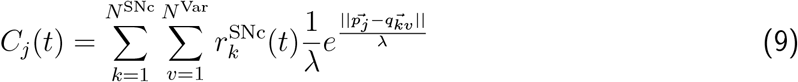

where 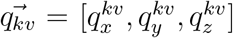 is the position of the *ν*th varicosity of SNc neuron *k*. λ controls the rate of decay with distance and was set to 0.1 (so that dopamine concentration decreases to 1/*e* ≈ 37% after diffusing a distance equivalent to 10% of the side of the cube). Under this model, there is an effective nigrostrital weight

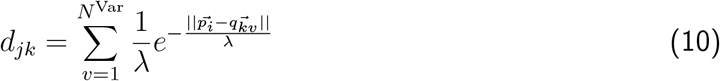

that does not vary with time, so we can write

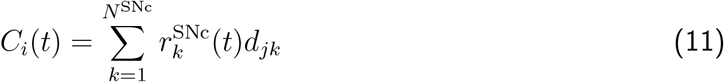

Note the similarity between Eqs. 11 and 7. If we introduce plasticity in the striatofugal projection, feedback alignment will cause 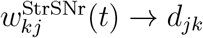. Therefore, we again start with

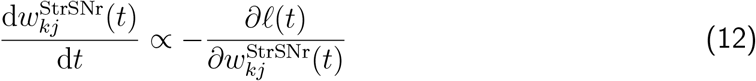

and arrive at

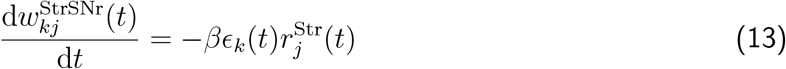

with 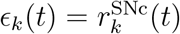 as before and *β* =10^−3^.

In summary, we set the corticostriatal plasticity update to

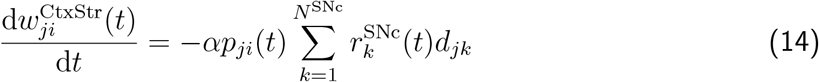

where *d_jk_* is given by Eq. 10 and *p_ji_*(*t*) is given by Eq. 6. We set *α* = –2.5·10^−2^ for direct pathway neurons and *α* = 2.5·10^−2^ for indirect pathway neurons to capture the different effects of D1 and D2 receptors. The minus sign helps 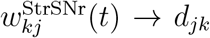 for direct pathway striatal neurons, because for these neurons 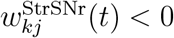. We used analogous plasticity rules for thalamostriatal and striatostriatal connections.

### Learning with vector valued dopamine feedback

Having set up the model, we simulated the dynamics for 5000 presentations of the target output (Fig. 1H). Each target was a 4-dimensional 200ms time series drawn from a Gaussian process (Fig. 1I; see Methods). In the first trial, the output does not match the target (Fig. 1I, left), but after 5000 trials the plasticity rules have driven the network to produce GPi/SNr output that closely matches with the targets (Fig. 1I, right). This is achieved both by plasticity in the readout population (GPi/SNr; Eq. 14) and plasticity in the striatum that adapts the SPN firing rates (Eq. 14; Fig. 1H).

We next asked how this learning depends on the nature of the dopamine feedback. It is well-known that for recurrent networks with rich dynamics, plasticity in the readout is sufficient to learn complex patterns (Jaeger and Haas, 2004; Maass and Markram, 2004). Therefore, we first compared our model to a reduced model that only had plasticity in the readout (striatofugal) projection according to Eq. 13, i.e. no plasticity in the striatum. We found that learning with dopamine feedback was faster (Fig. 1G). Next, we compared our model to a model in which each SPN at every timepoint receives the same dopamine feedback, i.e. the striatum receives a homogeneous, scalar dopamine signal. Strikingly, this model performs no better than the reduced model that only has learning in the readout layer (Fig. 1G). The increase in learning performance persisted with different sizes of the striatal (Supplementary Fig. S1A) and the readout (Supplementary Fig. S1B) populations, as well as for faster and slower timescales of the target (*τ*_task_ in Eq. 19; Supplementary Fig. S1C). These observations show that vector-valued dopamine feedback is crucial for the improvement in learning.

### The improvement in learning is because of feedback alignment

We next wanted to further explore the conditions during which vector-valued feedback improves learning of the targets. First, we considered two additional alternative models: (i) the feedback is random, i.e. we still have vector-valued feedback, but shuffle the coefficients *d_jk_* of Eq. 11 and (ii) we use the “ideal” feedback 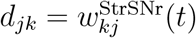. Note that locking the feedback weights to the feedforward in the second model means the feedback matrix is time-dependent. Both of these models perform similarly as our dopamine model (Fig. 2A). This suggests that the main criterion for the feedback to be effective is that feedback matrix *D* = [*d_jk_*] is non-degenerate.

**Figure 2:**
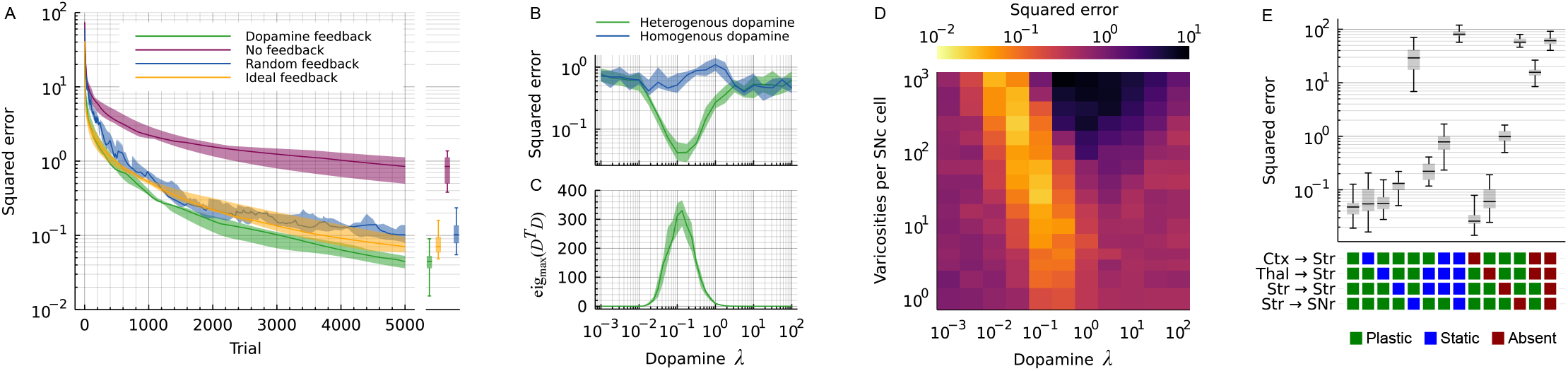
Learning improvement depends on feedback alignment. (A) Convergence of the loss (L2) for different types of feedback: dopamine feedback as described by our model, no feedback at all (effectively making all synapses except the striatofugal static), random feedback where the elements of the feedback matrix *D* are shuffled randomly and ideal feedback where 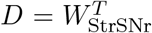. (B) Squared error after 5000 trials when some of the four plastic projections is impaired: either set to *static*, i.e. no plasticity, or removed from the model altogether (*absent*). (C) Squared error after 5000 trials as a function of the spatial constant of dopamine diffusion (λ). λ is measured in relation to the side of the cube in Fig. 1D. (D) Linear independence of the rows of *D* as a function of the dopamine spatial constant λ. (E) Squared loss after 5000 trials as a joint function of the dopamine spatial constant λ and the number of varicosities per SNc cell. See Supplementary Fig. S1D for the complete set of manipulations. All panels) errors are reported as the median over 25 trials. Shaded areas indicate the first and third quartiles of the 25 trials. Boxplots show median, first and third quartiles; whiskers indicate min and max.

With this hypothesis in mind, we tested varying the spatial scale λ of dopamine diffusion (see Eq. 10). Note that this spatial scale is measured as a fraction of the side of the cube (Fig. 1D). When λ ≪ 1, almost no dopamine reaches any SPN from the varicosities, and the network reverts to the *No feedback* control model (Fig. 2B). On the other hand, when λ ≫ 1, the dopamine from each varicosity covers the entire cube so that all SPNs effectively receive the sum of the dopamine released anywhere. This causes the network to revert to the *Homogeneous dopamine* control model. In between these two extremes, where dopamine scale is intermediate, there is a sweet spot where each SPN receives dopamine corresponding to a unique random linear projection of the 4-dimensional error (Fig. 2C). In this regime, the benefit of the feedback is the largest (Fig. 2B). Finally, we also tried to vary the number of varicosities *N*^Var^ in Eq. 10 and found that with a larger number *N*^Var^, a smaller spatial scale λ becomes viable (Fig. 2D). This is also consistent with the creation of a non-degenerate *D*.

To illustrate the importance of striatofugal plasticity for learning, we simulated the network model with “lesioned” basal ganglia projections by either removing the plasticity (*static*) or clamping them to 0 (*absent*). As expected, when the plasticity of the striatofugal projection was turned off, no feedback alignment could take place and the striatal plasticity could not contribute to learning the targets (Fig. 2E, Str→SNr). Turning off plasticity of the striatal projections (corticostriatal, thalamostriatal, and striatostriatal) on the other hand has a more moderate impact. This is because even with all of them fixed, we can still have echo-state-like learning in the striatofugal weights (see the *No feedback* null model in Fig. 1G and 2A). Similarly as when fixing the weights, removing either the cortical or the thalamic projection does not change the eventual error much, as both projections play similar and mostly interchangeable roles in our model, whereas removing both silences the striatum completely and hence gives a very large error (Fig. 2E). See Supplementary Fig. S1D for a systematic investigation how removing or blocking plasticity on different projections affect learning.

### Fast synaptic dynamics can compensate for slow dopamine

So far we have assumed that dopamine is diffuse in space, but delivered instantly to the receiving SPNs. While this allows the synaptic weights to be updated correctly on every time step, it neglects the temporal dynamics of dopamine release, diffusion, reuptake, etc. A faithful quantitative model of these processes is beyond the scope of our abstract rate network, but it is nevertheless important to determine how dependent our dopamine-based learning rule is on the assumption of instantaneous dopamine release. We simplified all temporal dynamics of dopamine into a simple exponential low-pass filter with time constant *τ*_DA_. That is, we changed the equation for dopamine concentration at striatal neuron *j* (Eq. 9) to

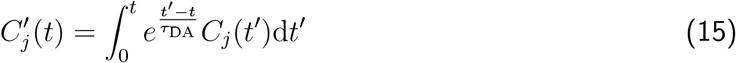

The blue line in Fig. 3A shows the resulting error after 5000 trials for a range of values of *τ*_DA_. When *τ*_DA_ is much faster than the time constant of the task (here *τ*_task_ = 20ms), the error (Fig.3A, blue line) is similar to the earlier, instantaneous dopamine model (Fig.3A, green line). However, for slower *τ*_DA_ the error increases and even surpasses the null model with no striatal plasticity (Fig.3A, purple line).

**Figure 3:**
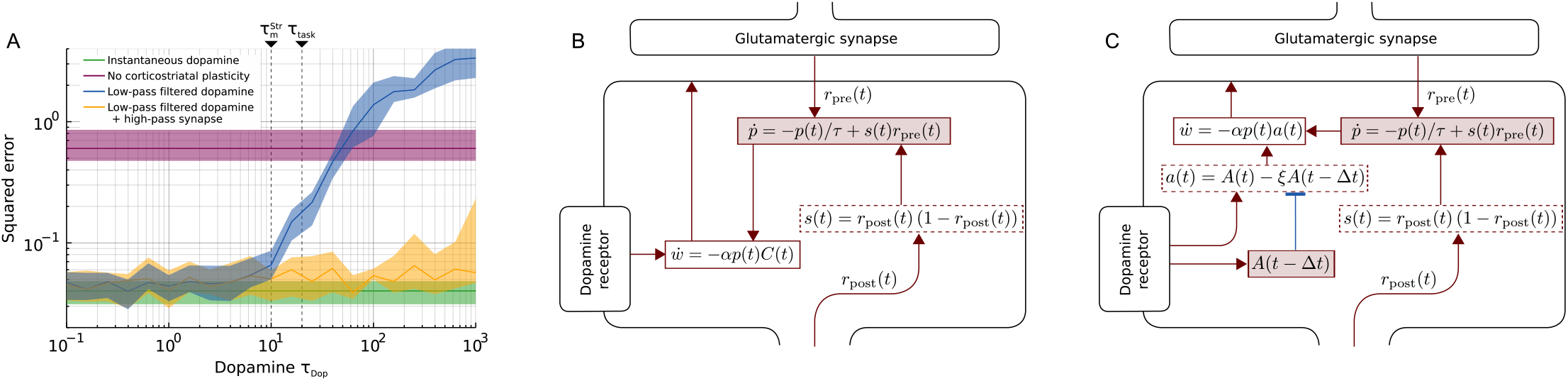
Slow dopamine dynamics. (A) When the dopamine signal is low-pass filtered with a time constant slower than the time constant of the target output, the error (blue) increases. For comparison, the errors of previous models with instantaneous (green) and absent (purple) dopamine are also shown (these do not depend on the *τ*_Dop_). Finally, the orange line shows the error when using the extended synaptic dynamics shown in panel C. All errors are reported as the median over 25 trials. Shaded areas indicate first and third quartiles of the 25 trials. (B) The synaptic dynamics of the standard RFLO learning rule. (C) Adding an extra forward-inhibition motif to the synaptic dynamics to compensate for temporal smoothing of dopamine.

We next asked what would be needed to rescue the learning performance in face of slow dopamine dynamics. One possible solution would be that each synapse high-pass filters its local dopamine concentration. Such high-pass filtering can be done by assuming a feed-forward inhibition motif as the first step in the biochemical pathway triggered by dopamine (Fig.3C). In the ideal case one biochemical node in the each synapse is tracking dopamine concentration one time step ago, and another calculates the difference

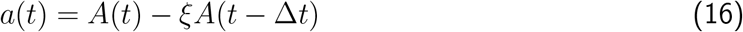

Furthermore, if we choose 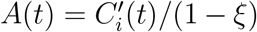 and 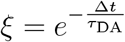 we get *a*(*t*) ≈ *C_i_*(*t*) and we can use the same RFLO learning rule as before but with *a*(*t*) instead of *C_i_*(*t*) (compare Figs 3B and 3C).

We verified this idea by introducing the synapse model in Fig.3C in all striatal synapses and then again plotting the error after 5000 trials. As predicted by the ideal choice of *ξ*, the error was consistently similar to instantaneous even for large *τ*_DA_ (Fig.3A, yellow line).

## Discussion

The broad, unspecific dopaminergic axonal projections have been argued to only allow for the transmission of a scalar homogeneous feedback signal (Chen and Goldberg, 2020). Here we provide a tenable counter-example of this view, even if speculative and highly idealized. We demonstrate that in a reduced model of a piece of basal ganglia that a heterogeneous, vector-valued feedback signal could in fact be transmitted by dopamine, even if the dopaminergic projections in the striatum are random. We have identified four key requirements for effective use of vector-valued dopamine, which also serve as predictions that can be verified experimentally:

1. At least one projection downstream of the striatum must be plastic.
2. The vector-valued error must be available to both the dopaminergic (here, SNc) and the readout (here, SNr) populations.
3. Striatal dopamine dynamics must be at least as fast as the targeted movements, or, alternatively, be high-pass filtered by feed-forward inhibition in the synaptic biochemical pathways.
4. The dopamine received by each SPN must be sufficiently independent, or, stated formally, the effective connectivity matrix arising from summing contribution of individual varicosities (Eq. 10) must not be degenerate.

Direct experimental evidence for any downstream plasticity in the basal ganglia (*Requirement 1*) is scarce, but in a recent study González-Rodríguez et al. (2021) showed that dopamine depletion in the SNr plays a larger role than striatal dopamine in producing motor deficits in Parkinson’s disease. This is qualitatively consistent with the relative importance of striatonigral over corticostriatal plasticity in our model (Fig. 2F). However, we note that although we placed this plasticity in the striatofugal projection(s), it could in principle also be met by plasticity in the nigrothalamic or nigrocollicular projections.

A vector-valued error would likely appear as tunings to various motor and task variables in experimental animals (*Requirement 2*), especially in the phase before the animals are so overtrained their error is zero. Indeed, cells in the SNr (Fan et al., 2012; Barter et al., 2015; Tang et al., 2021) as well as the SNc (Howe and Dombeck, 2016; Dodson et al., 2016; Coddington and Dudman, 2018; Avvisati et al., 2022) respond to a plethora of behavioral and task variables. We deliberately excluded the details of how this error may be computed in the brain, but we speculate at least three possible algorithmic ways in which it could appear:

1. Bran region regions such as motor cortex or cerebellum could have a forward model of the world as well as the target, and thus, can directly compute the error and send it to the midbrain.
2. The brain could be wired as a set of hierarchical control loops, in which each loop provides the target for the level below (as proposed by Yin, 2014). Each such loop could stretch throughout the cortex and basal ganglia.
3. If the executed action has more variability than the command read out by the SNr, the *policy gradient theorem* (Sutton et al., 1999) states that the gradient for the update should be

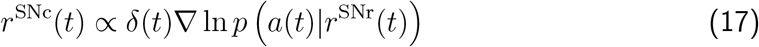

where *α* is the vector-valued action taken, *δ* is the temporal difference (TD) error as predicted by a critic, and *p* is the probability density function of *α*. Note that this suggests that SNc cells should fire proportionally to both the TD error and to (the gradient of) some behavioural variables, which could explain why many SNc cells appear tuned to both (Parker et al., 2016; Lee et al., 2019).Lindsey and Litwin-Kumar (2022) have proposed dorsal striatum could make use of such a policy gradient, but nonetheless argue dopamine itself is a scalar proportional to the squared norm of the policy gradient.

There are several ways the brain could implement filters that allow extraction of faster fluctuations in dopamine concentration (*Requirement 3*). Our example in Fig. 3 is highly idealized and makes arguably unfair use of our idealized perfect exponential decay of the dopamine. In reality, the journey of dopamine molecule from a varicosity to a dopamine receptor depends on the local geometry and dopamine reuptake so that the dependence on both time and distance is most likely complicated and non-linear (although these effects might less pronounced at very short distances; Cragg and Rice, 2004; Liu et al., 2021). Nevertheless, evolution has had a good opportunity to tweak the biochemical pathways to compensate for these effects as far as permitted by the signal-to-noise ratio. Whether this is tenable in a realistic model of dopamine diffusion and biochemical cascades remains an open question, but we predict that there is at least one node in the biochemical cascade of dopamine-induced synaptic plasticity that is sensitive to fast fluctuations in local dopamine concentration.

The spatial frequency of the dopamine landscape in the striatum must be high enough so that even neighbouring SPNs do not sense the exact same dopamine concentration (*Requirement 4*). This can be achieved by having a short spatial constant of dopamine diffusion, and possibly compensating with a larger number of varicosities (Fig. 2D). Consistent with our model, Cragg and Rice (2004) estimated the diffusion distance of dopamine following release to a few microns.

The main goal of our work was to demonstrate that the broad and unspecific nigrostriatal dopaminergic projection can in principle transfer a usable vector-valued error to the striatum; our ambition was not to provide a complete biological account of the process. For this reason, there are many likely very important features of basal ganglia anatomy and physiology we did not include, for example dorsolateral/dorsomedial functional division in the striatum (Balleine and O’Doherty, 2010), the different roles of the matrix and the striosome (Bloem et al., 2017), axonally initiated dopaminergic release by cholinergic interneurons (Threlfell et al., 2012; Liu et al., 2022), saturating dopamine receptors (Liu et al., 2021), etc. Similarly, our primary goal was not to introduce a new algorithm for training recurrent neural networks; the network setup and plasticity rule is an application of the RFLO rule (Murray, 2019). Nevertheless, we show that the RFLO rule is applicable in a basal ganglia-like network with multiple inhibitory synapses and with our toy model of dopamine feedback, and propose vector-valued error feedback as a candidate functional role of dopamine.

Previous proposals for use of heterogeneous dopamine (Hamid et al., 2021; Lee et al., 2022) assume that the heterogeneous responses of dopaminergic cells are transmitted to the striatum through private parallel channels without any cross-talk. However, this not easily reconciled with functional and anatomical findings (see Introduction). Another proposed use of heterogeneous firing in the midbrain dopaminergic neurons is to support a distributional coding of value (Dabney et al., 2020). However, a distributional value code only explains different gains in the coding of the reward prediction error, not why the neurons respond to non-rewarded task variables. Nevertheless, it is entirely possible that the brain simultaneously employs a distributional value code (perhaps most strongly in the VTA) and a vector-valued error code (perhaps most strongly in the SNc).

In conclusion, we propose that the heterogeneous responses of dopamine cells seen by Fan et al. (2012); Howe and Dombeck (2016); Engelhard et al. (2019) and others represent a vector-valued error. By providing this type of error, the SNc supports the basal ganglia learning to select actions from a continuous action space in continuous time, thereby providing the animal with vital behavioral flexibility, control and adaptability.

## Acknowledgements

We thank Dr. Moritz Weglage and Prof. Erik Fransén for helpful discussions. Partial funding from StratNeuro (to AK), Swedish Research Council (to AK) and Digital Futures (to AK) and Karolinska Institutet (KID doctoral funding to EW) is gratefully acknowledged.

## Methods

The dynamics of neurons, network structure and learning rule is already described in the Results section. Here we describe only the technical details needed to run the simulations.

### Network simulations

The network was simulated in a custom simulator written in Julia (Bezanson et al., 2017). The dynamics were simulated with forward-Euler with dt=1ms.

Simulations consisted of multiple trials concatenated after each other without any reset of the network in between. The current time in the current trial was signaled to the network by setting the thalamic firing rates to

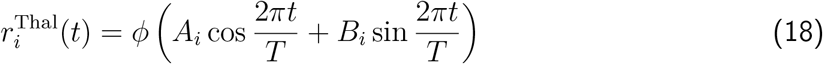

where *T* = 200ms is the duration of a single trial, and *A_i_* and *B_i_* are constants drawn randomly from a circle with radius 4 (i.e. 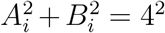 for all *i*). *ϕ* is the logistic transfer function (Eq. 2).

### Initializing the weights

For each pair of cells in each projection, there was a fixed probability (Fig. 1B) of a synapse being inserted. If a synapse was inserted, its weight was drawn from a uniform distribution 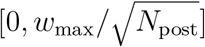 (see Table S1) and then multiplied by −1 for the inhibitory projections. The weights in Table S1 were chosen for the network to have close to chaotic trajectories before training.

After all synapses were created, the sum of the weight of all incoming synapses was calculated for each neuron. If this sum was greater than 0, all the inhibitory synapses were slightly increased so that the new sum was exactly 0. Conversely, if the sum was less than 0, all the excitatory weights were slightly increased to reach sum 0. This ensured that each neuron had roughly balanced excitation and inhibition, which in turn created rich dynamics from the start.

### Target signals

The targets were drawn from a Gaussian process with mean 0.5 and variance given by

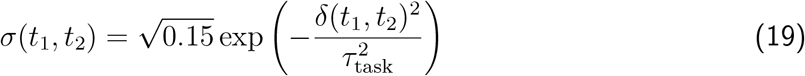

where *δ*(*t*_1_, *t*_2_) is the smallest difference between *t*_1_ and *t*_2_ when including wrap-around, i.e.

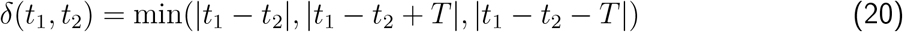

where *T* = 200ms is the duration of a single trial. The periodic kernel is to avoid discontinuities when running consecutive trials without resetting the network. For all experiments in the main figures, *τ*_task_ = 20ms.

**Table S1:** Initial synaptic strength. Note that weights for inhibitory connections were multiplied by −1 after they are drawn.

| pre | post | $w_{\max}$ |
| --- | --- | --- |
| Cortex (exc) | Cortex(exc) | 25 |
| Cortex (exc) | Cortex(inh) | 25 |
| Cortex (inh) | Cortex(exc) | 25 |
| Cortex (inh) | Cortex(inh) | 25 |
| Cortex (exc) | Striatum (dSPN) | 25 |
| Cortex (exc) | Striatum (iSPN) | 25 |
| Striatum (dSPN) | Striatum (dSPN) | 25 |
| Striatum (dSPN) | Striatum (iSPN) | 25 |
| Striatum (iSPN) | Striatum (dSPN) | 25 |
| Striatum (iSPN) | Striatum (iSPN) | 25 |
| Striatum (dSPN) | SNr/GPi | 5 |
| Striatum (iSPN) | SNr/GPi | 5 |
| Thalamus | Cortex (exc) | 50 |
| Thalamus | Cortex (inh) | 50 |
| Thalamus | Striatum (dSPN) | 30 |
| Thalamus | Striatum (iSPN) | 30 |

**Figure S1:**
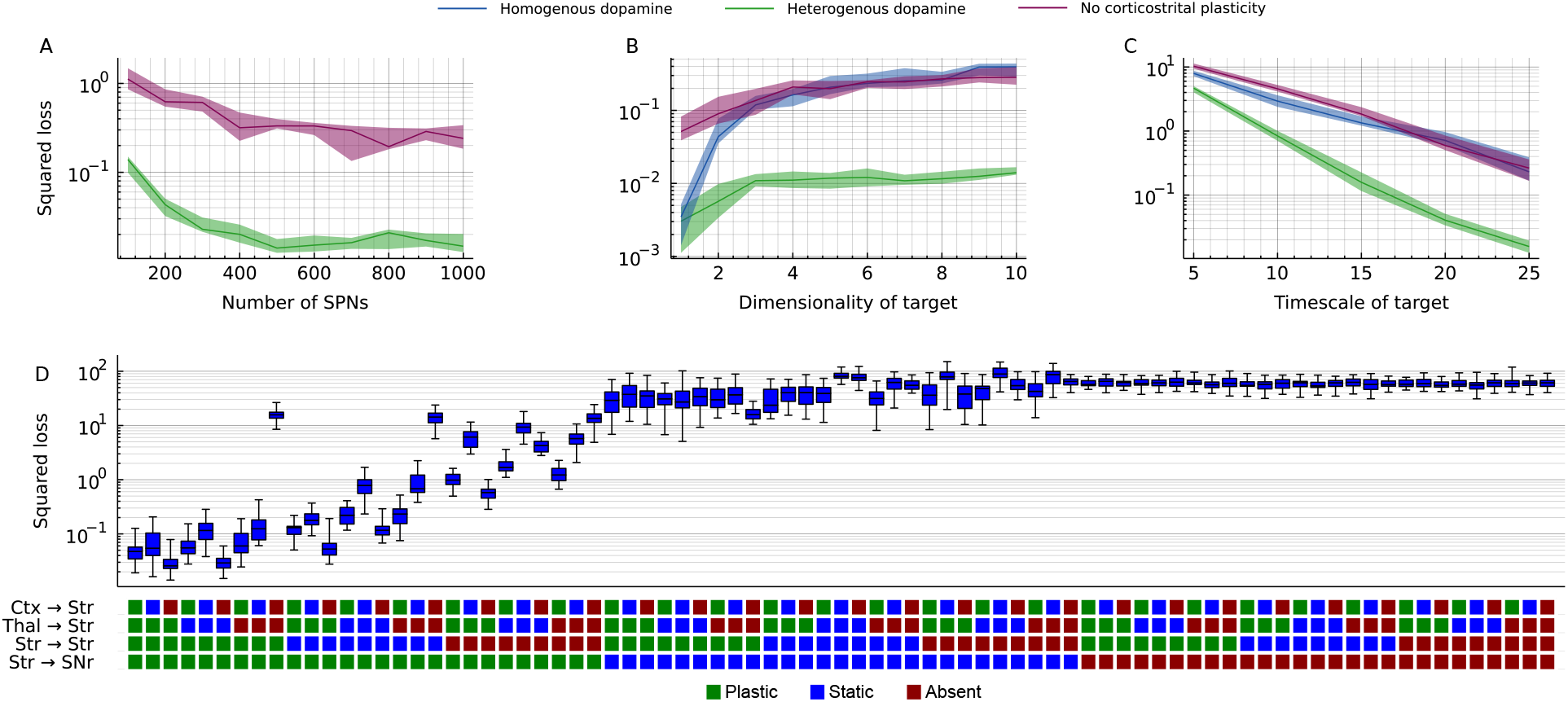
Influence of each projection to the loss. (A) Mean squared error after 5000 training trials for different sizes of the striatal population. (B) Mean squared error for increasing values of *N*^SNr^ = *N*_SNc_ = *d*. (C) Mean squared error for increasing values of *τ*_task_. (D) The network model was run with impairments to some of the connections. Either the plasticity was removed so that the synaptic weights of the connection were fixed to their starting weights (“static”), or the synapses were removed altogether (“absent”). Squared error is measured as the error after 5000 trials. Boxplots show median and quartiles across 25 runs; whiskers indicate min and max of the 25 runs.

### Derivation of the plasticity rule

We begin by restating the network equations. First, we write out the three inputs (cortex, thalamus and recurrent striatum) as

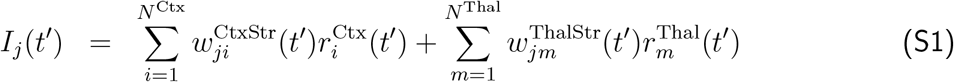

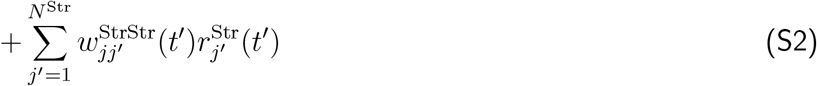

Then we rewrite the membrane equation as an integral

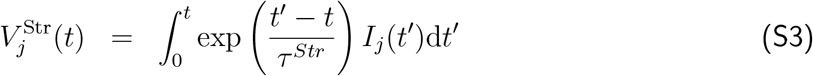

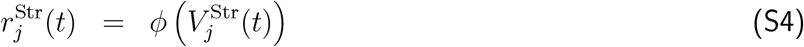

Since we treat SNr as a read-out layer, we let *τ*^SNr^ → 0 so that

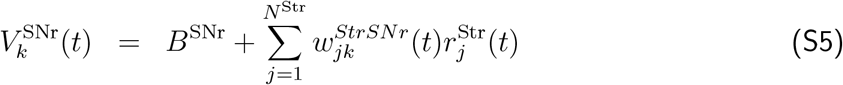

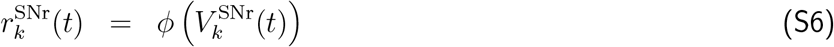

The instantaneous loss is

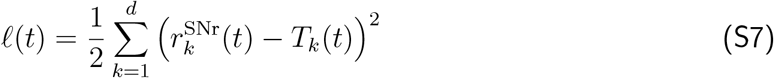

where 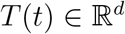 is the target output at time *t*. To greedily (i.e. not considering earlier or later losses) minimize *ℓ*(*t*) (see Murray, 2019) we want to have

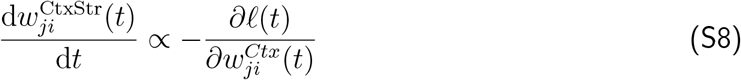

Expanding this taking a partial derivative where *r*_*j*′_(*t*) is fixed for *j*′ ≠ *j*:

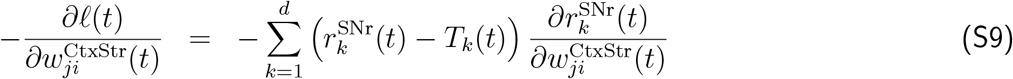

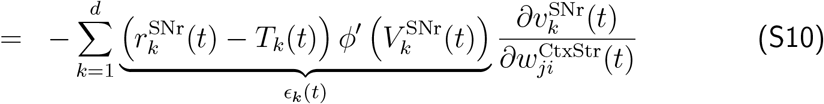

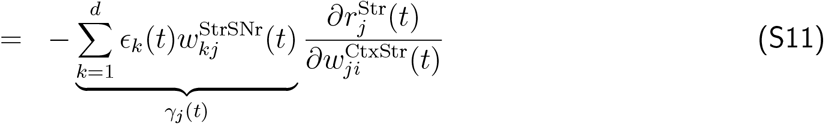

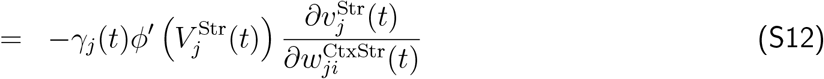

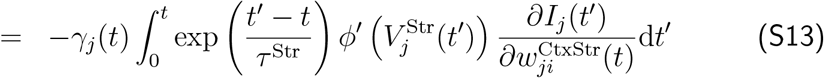

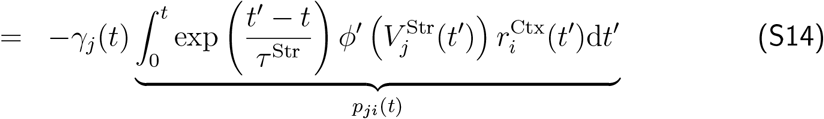

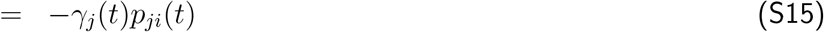

Furthermore, note that

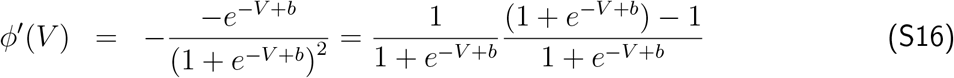

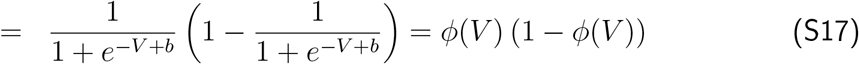

In summary, we get the following update rule for the corticostriatal synaptic weights

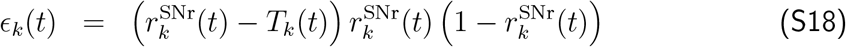

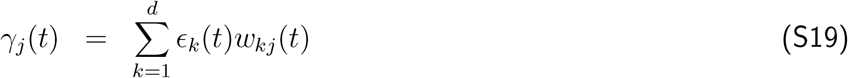

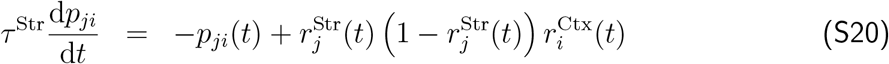

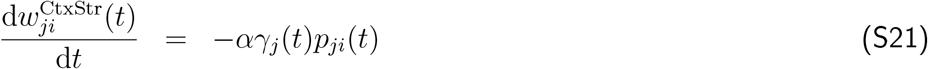

The update rules for thalamostriatal (*w*^ThalStr^) and striatostriatal (*w*^StrStr^) have the same form and are derived in the same way. The striatofugal weight plasticity is

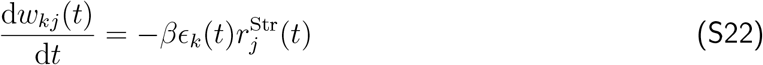

with the same *ε_k_* as above.

